# Molecular programs of fibrotic change in aging human lung

**DOI:** 10.1101/2021.01.18.427195

**Authors:** Jasmine Lee, Mohammad Naimul Islam, Kaveh Boostanpour, Dvir Aran, Stephanie Christenson, Michael A. Matthay, Walter Eckalbar, Daryle J. DePianto, Joseph R. Arron, Liam Magee, Sunita Bhattacharya, Rei Matsumoto, Masaru Kubota, Donna L. Farber, Jahar Bhattacharya, Paul J. Wolters, Mallar Bhattacharya

**Author notes:** Equal contribution (Jasmine Lee, Mohammad Naimul Islam, Jahar Bhattacharya, Paul J. Wolters, Mallar Bhattacharya).

## Abstract

Aging is associated with both overt and subclinical lung fibrosis, which increases risk for mortality from viruses and other respiratory pathogens. The molecular programs that induce fibrosis in the aging lung are not well understood. To overcome this knowledge gap, we undertook multimodal profiling of distal lung samples from healthy human donors across the lifespan. Telomere shortening, a cause of cell senescence and fibrosis, was progressive with age in a sample of 86 lungs and was associated with foci of DNA damage. Bulk RNA sequencing confirmed activation of cellular senescence and pro-fibrotic pathways as well as genes necessary for collagen processing with increasing age. These findings were validated in independent datasets for lung and sun-exposed skin, but not other organs including heart, liver and kidney. Cell type deconvolution analysis revealed a progressive loss of lung epithelial cells and an increasing proportion of fibroblasts. Consistent with the observed pro-fibrotic transcriptional profile, second harmonic imaging demonstrated increased density of interstitial collagen in aged human lungs. Furthermore, regions of parenchymal fibrosis were associated with decreased alveolar expansion and surfactant secretion. These findings reveal the transcriptional and structural features of fibrosis and associated physiologic impairments in normal lung aging.

## Main

Lung capacity and resilience decline and susceptibility to disease increase with age^1^, but the molecular and structural mediators of this natural history are unknown^2^. Targeting respiratory aging therapeutically or prophylactically will require understanding of lung-specific molecular programs that change with age^3-9^. To characterize the effect of age on gene expression in lung, we prospectively collected 86 human donor lungs as part of the Lung Aging Cohort (LAC).

Lungs were evenly distributed in age between 16 and 76 years (**Figure 1a-b**). Donors were not known to have any underlying pulmonary conditions, and gender, smoking status, and ethnicity are summarized in **Figure 1c** and detailed in **Table S1**. Tissue samples were harvested from distal lung and frozen in liquid nitrogen on receipt. RNA was later extracted from these samples for Illumina-based sequencing in a single run after cDNA library preparation.

**Figure 1.**
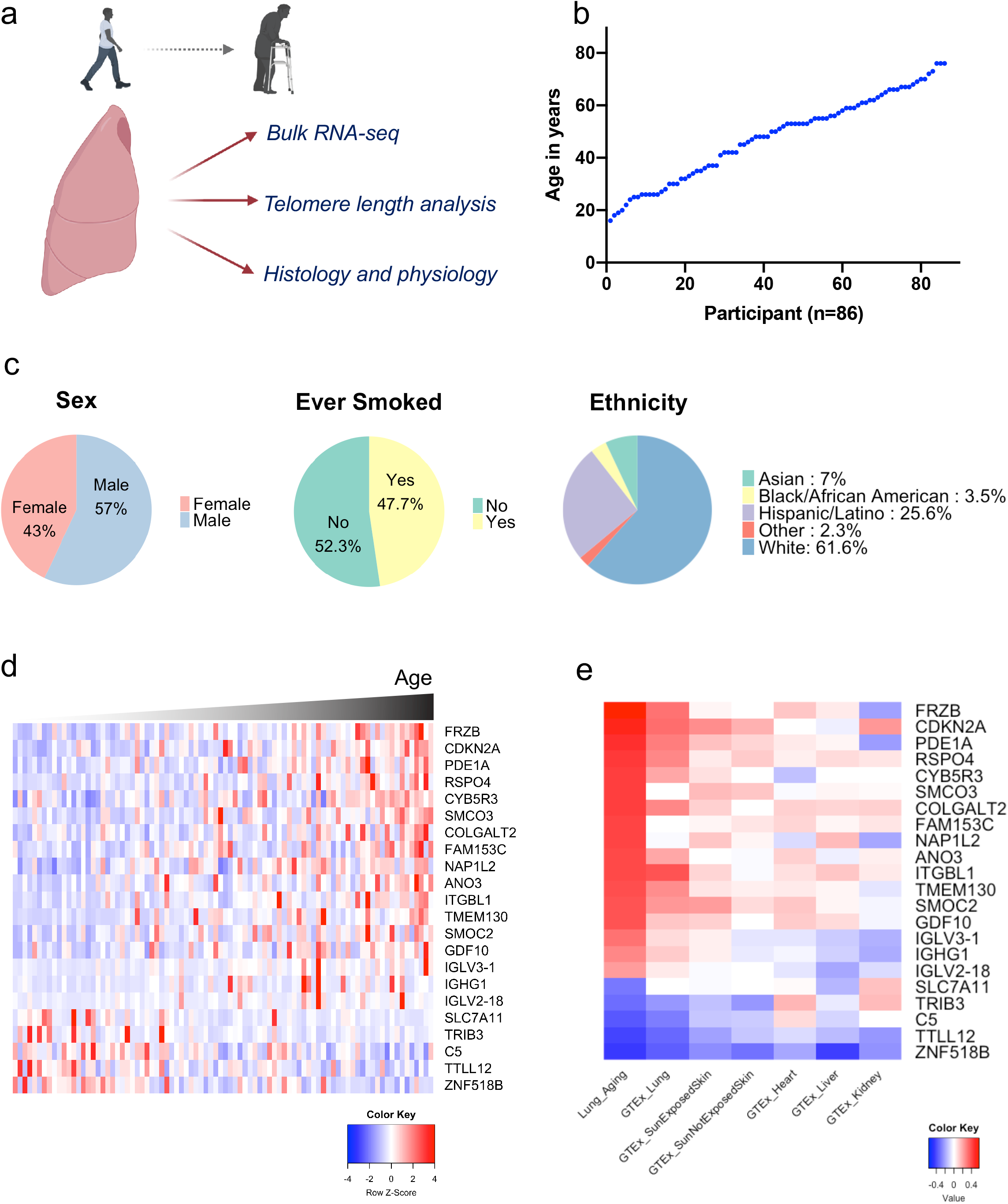
RNA-seq reveals a lung-specific aging signature. **a**, Schematic of the lung aging cohort (LAC), a study of human lungs varying in age across the adult lifespan and profiled with multiple approaches. **b**, Participant age plotted against increasing order of age. **c**, Demographic features. **d**, Heatmap of gene expression by bulk RNA-seq of distal lung. Genes listed were significantly correlated with age in a multivariate generalized linear model controlling for smoking and gender (FDR p<0.1). Z scores represent within-gene relative expression across samples. **e**, Heatmap of Pearson correlation coefficients between gene expression and age for the LAC and multiple GTex tissue datasets.

Differential gene expression analysis using a multivariate linear model controlling for gender and smoking identified 22 genes that correlated significantly with age as a continuous variable in the LAC (**Figure 1d**). To validate this lung aging gene signature, we used publicly available data from the Genotype-Tissue Expression (GTEx) project of multiple tissues from over 300 individuals^10^. The LAC gene signature was also associated with age in the GTEx lung samples. Interestingly, overlap of the lung aging gene signature was found for aging of sun-exposed regions of skin, but not for non-sun-exposed skin, kidney or heart (**Figure 1e**).

Given the relevance of cellular senescence to aging, samples were interrogated for evidence of cell senescence markers and pathways. The canonical senescence marker p16 (CDKN2A) was among the most highly upregulated genes in aging lungs (**Figure 2a**). To further assess for senescence reprogramming, we asked whether consensus senescence gene signatures that we recently defined by RNA-seq of senescent lung epithelial cells and fibroblasts^11^ were upregulated in the LAC. The consensus senescence gene signatures were increased with age in both the LAC and GTex Lung datasets (**Figure 2b**). We then performed Ingenuity Pathways Analysis (IPA) of genes associated with aging in the LAC (**Figure 2c**; **Table S2**), identifying pathways that were largely validated in GTex lung (**Figure 2d**). Consistent with cellular senescence, cell proliferation pathways were inhibited, and both p16 (CDKN2A) and p21 (CDKN1A) pathways were activated in aged lungs. Cell death pathways were also prominently activated. Overall, the marker-based analysis and IPA suggest activation of senescence and cell stress in aged lungs at steady state.

**Figure 2.**
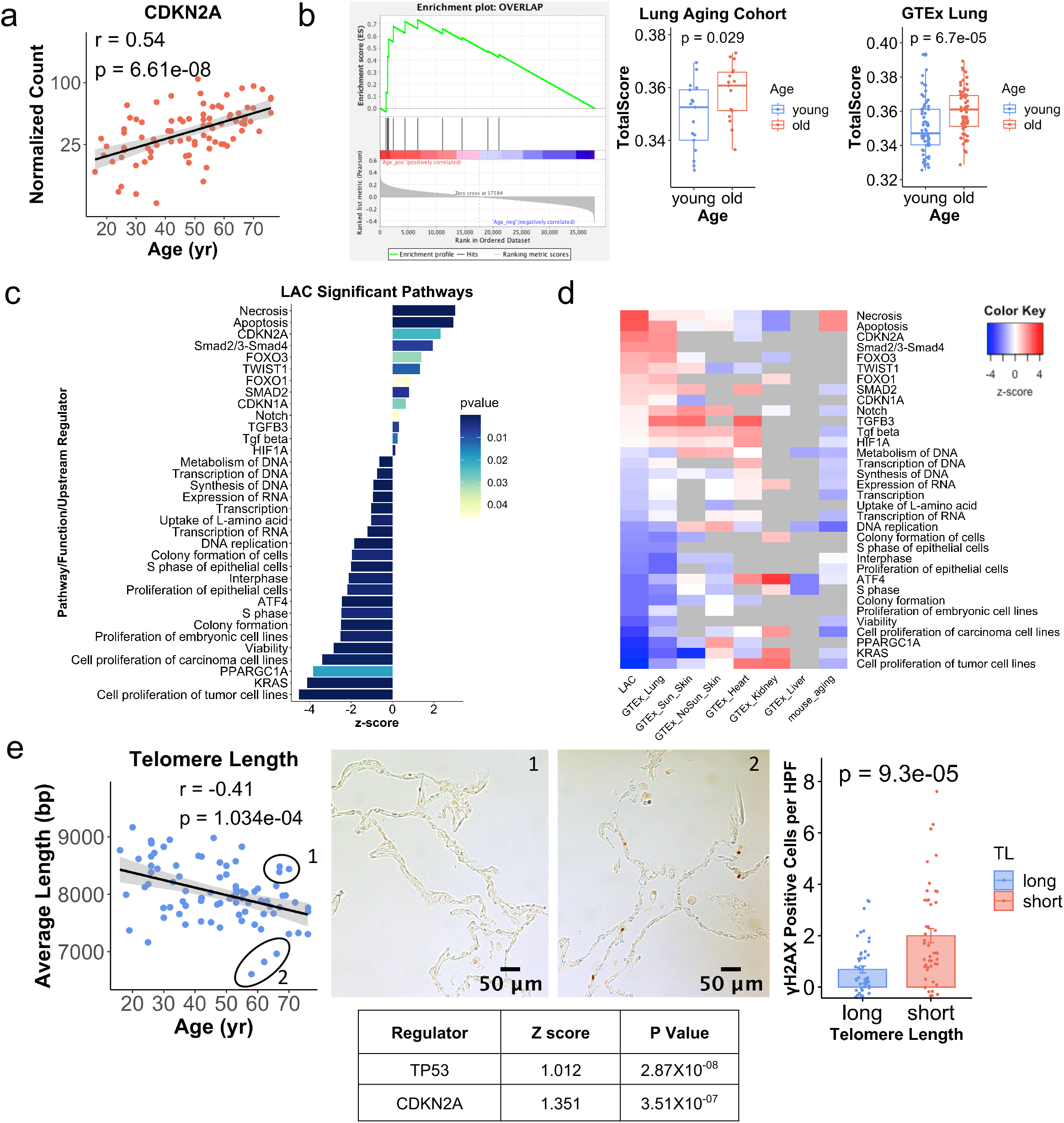
Lung cellular senescence increases with aging. **a**, p16 expression by age in the LAC (log scale). Pearson R is shown. **b**, Gene set enrichment analysis of 11 consensus senescence markers from DePianto et al11 (p=0.004). Single sample geneset enrichment analysis using the 11-gene signature is shown for individuals in the LAC (total N=34) and GTex lung (total N=138). P values are for two-sided Student’s t-test. **c**, Ingenuity Pathways Analysis (IPA) computed with the genes from LAC that were correlated with age at p<0.05 level of significance. **d**, Genes used for IPA in (c) were used to compute IPA results for the same pathways in multiple other datasets. Z-scores are shown for pathways reaching significance at P<0.05. Pathways in grey were not detected or not significant. **e**, qPCR-quantified telomere length plotted by age for the LAC. Pearson R is shown. The circled subsamples differ in telomere length (1=long, 2=short) and were used for gamma-H2ax immunohistochemistry, with representative images and quantitation to the right (n=3 in each group, two-tailed Student’s t-test p value is shown). Differential gene expression of the subsamples by bulk RNA-seq was used for IPA and predicted activity of upstream senescence regulators in the short telomere subgroup, with Z scores and p values shown in the table.

Cellular senescence has been associated with fibrotic lung disease, in part due to a senescence-associated secretory profile that has pro-fibrotic effects^12-18^. Therefore, we considered that cell senescence may also underlie aging-associated subclinical interstitial fibrosis, or interstitial lung abnormalities, a phenomenon recognized recently by radiographic studies of asymptomatic aged individuals^19-21^. These radiographic findings of fibrosis have been correlated with histopathologic features of fibrosis, including fibroblastic foci and subpleural distribution^22^. However, little is known about the molecular and cellular programs responsible for the pro-fibrotic evolution in aging lung. We first noted that pathways consistent with mesenchymal activation and fibrosis (TGF-beta pathway mediators and the epithelial-to-mesenchymal transition regulator TWIST1) were activated in aged lungs (**Figure 2c-d**). These results were largely confirmed in the GTEx lung dataset but not consistently in other organs (**Figure 2d**). Furthermore, several of the most highly upregulated genes (**Figure 1e**) have known pro-fibrotic effects; for example, RSPO4 has been associated with decline in lung function in patients with lung fibrosis^23^.

Next, since a major cell-intrinsic driver of senescence is telomere shortening, we isolated genomic DNA from the lung samples and used a quantitative PCR-based assay to measure telomere length. This analysis revealed that average lung telomere length progressively decreased across the lifespan (**Figure 2e**). Telomere attrition leads to telomere uncapping, which triggers a DNA damage response including p53 activation^24^. To test the significance of telomere shortening to cellular states, gene expression was compared between subsamples that were significantly different in telomere length but approximately matched by age. IPA upstream regulator analysis of differentially expressed genes revealed that the canonical senescence regulators p53 and p16 were activated in association with decreased telomere length; furthermore, sites of DNA damage were increased in the short-telomere samples by gamma-H2ax immunohistochemistry (**Figure 2e**). These results suggest that age-associated lung telomere attrition likely contributes to the senescence profile observed by IPA.

Given the senescence and cell death profiles revealed by our analysis, we next asked whether lung aging is associated with changes in the cellular composition of the lung. To address this question, cell type deconvolution analysis was performed on the bulk RNA-seq data. First, SingleR^25^ was used to annotate cell types from published single cell transcriptomes for 3 young, healthy human lungs.^26^ Differential gene expression analysis confirmed characteristic markers for lung epithelial cells and fibroblasts (**Tables S3** and **S4**, respectively). MuSiC^27^ was then applied to these SingleR-identified clusters of cell subtypes to deconvolve proportions of cell types in each LAC and GTex lung sample. Interestingly, the proportion of epithelial cells declined with age; on the other hand, the proportion of fibroblasts increased, consistent with fibrotic change in the aging lung (**Figure 3a; Figure S1a**). Within the epithelial compartment, we found specifically alveolar type 2 cells to decrease with age by immunostaining for the type 2 cell marker pro-SPC (**Figure 3b**). Type 2 cells are thought to be necessary for epithelial renewal in the lung, even at steady state.^28^ Collectively, these findings support a DNA damage response resulting from telomere shortening, leading to epithelial senescence and pro-fibrotic pathway activation characterized by expansion of the mesenchyme in the aging human lung.

**Figure 3.**
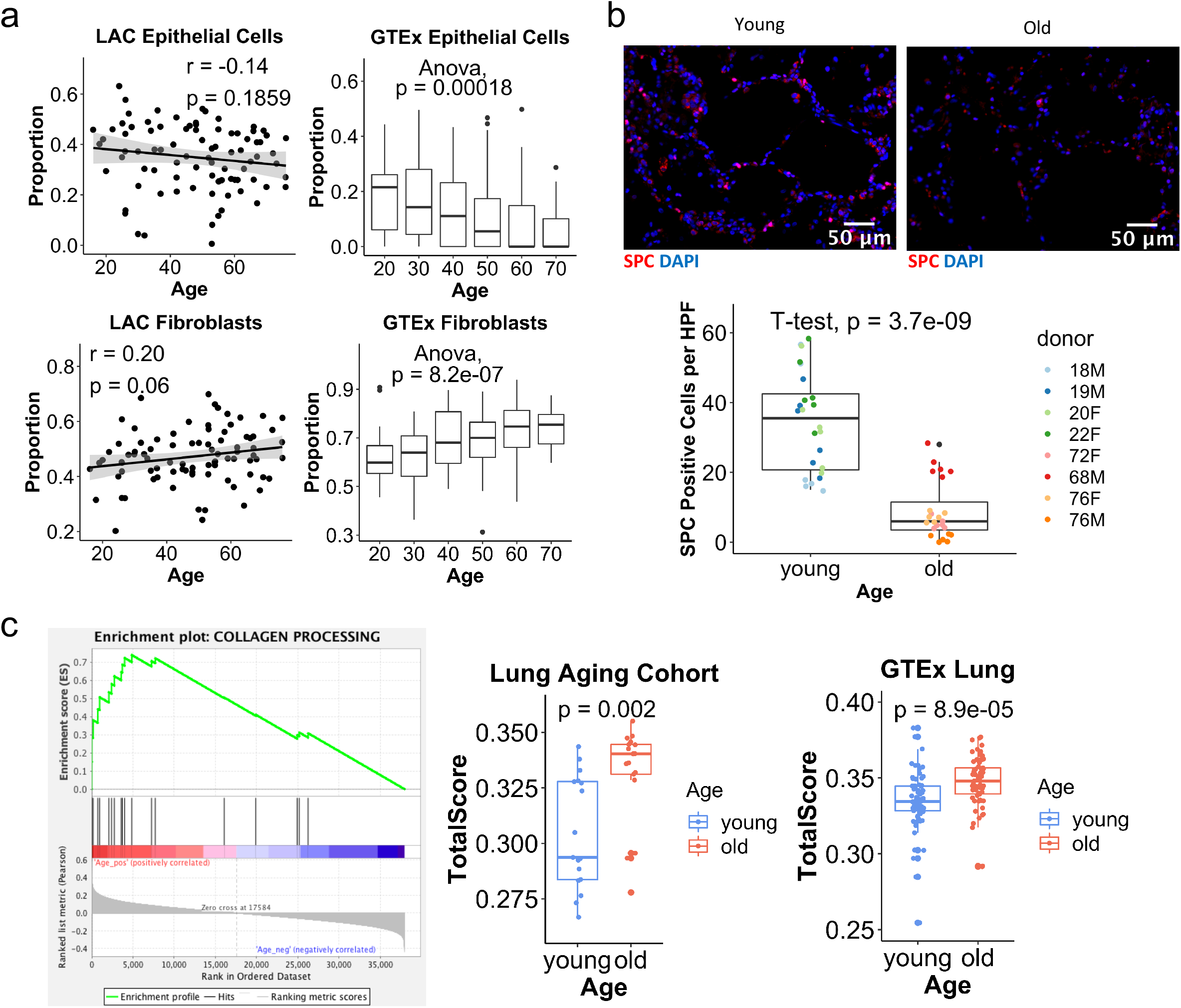
Fibrotic programs in lung aging. **a**, Cell type deconvolution of bulk RNA-seq data from the LAC and GTex Lung. Age for GTex is represented in decades. Pearson R and and p values are shown for LAC, and 1-way ANOVA is shown for GTex. Boxplots show the median, first and third quartiles, and 1.5^∗^IQR. **b**, Quantification of type 2 cells in old and young lungs by labeling of SPC+ cells, with quantification (N=4 individuals in each group, donor age and gender are shown). P value is for two-tailed Student’s t-test comparing all old and all young samples. **c**, Collagen-processing gene expression in the LAC by gene set enrichment analysis (p=2.004e-04). Single sample gene set enrichment analysis is plotted for the genes enriched in aging (genes listed in Fig S1) in the oldest and youngest quintiles from the LAC and GTex Lung. P values are for two-tailed Student’s t-test.

We noted no robust differences in expression of collagen genes between old and young. However, collagen accumulation is due not simply to excess collagen deposition, but also to an imbalance of collagen production and destruction, as well as changes in extracellular collagen structure and stability^29,30^. Therefore, the oldest and youngest quintiles in the LAC were examined for expression of genes known to regulate post-translational processing of collagen, including lysyl oxidases, transglutaminases, and tissue inhibitors of matrix metalloproteases, which inhibit collagen turnover by metalloproteases. Remarkably, a large proportion of these genes were upregulated with age (**Figure 3c; Figure S1b**). We then tested whether the age-associated subset of these genes from the LAC could be validated in other datasets and found robust upregulation of the signature in the GTex lung cohort (**Figure 3c**), and for GTex sun-exposed skin, but not for multiple other organs (**Figure S1c**).

These changes in gene expression and cellular content of the lung led us to test the age dependence of collagen structure and distribution in young and aged human donor lungs by live two-photon microscopy. Second harmonic (SH) generation by extracellular collagen has been used to visualize the fibrillary structure of collagen in fixed tissues.^31^ Here we applied the technique to live, unfixed human lungs. SH imaging revealed marked differences in the collagen pattern in young versus aged lungs in the subpleural space. In SH images of young lungs (age<40 years), well-defined collagen fibers of 1 micron thickness were regularly evident with interfibrillar spaces of 3-5 microns (**Figure 4ai**). Fluorescence analyses along lines drawn on the *x-y* planes of these images revealed a fluorescent spike where the analysis line intersected a fibril, while the low inter-spike fluorescence reflected the collagen-free interfibrillar space (**Figure 4b**). In aged lungs (age >65 years) a similar fibrillar pattern was also evident in some regions, but in other regions the spiked fibrillar pattern was notably absent (**Figure 4aii, 4b**). In these regions line analyses revealed dense packing of considerably thinner fibers (**Figure 4b**). Area analysis of SH fluorescence quantified in the Z direction starting at the pleural surface revealed Gaussian distributions of fluorescence intensity reflecting intensity and depth of subpleural collagen deposition (**Figure 4c**). Notably, subpleural collagen density varied considerably between different regions of the same lung for both young and old donors, as indicated by the spread of density values for each lung (**Figure 4d**). On average collagen density was higher in the older age group (**Figure 4e**). In the alveolar interstitium subjacent to the pleural space, collagen density was about ten times less than in the subpleural region (**Figure 4f**). However, here too older lungs had higher interstitial collagen density (**Figure 4g**).

**Figure 4.**
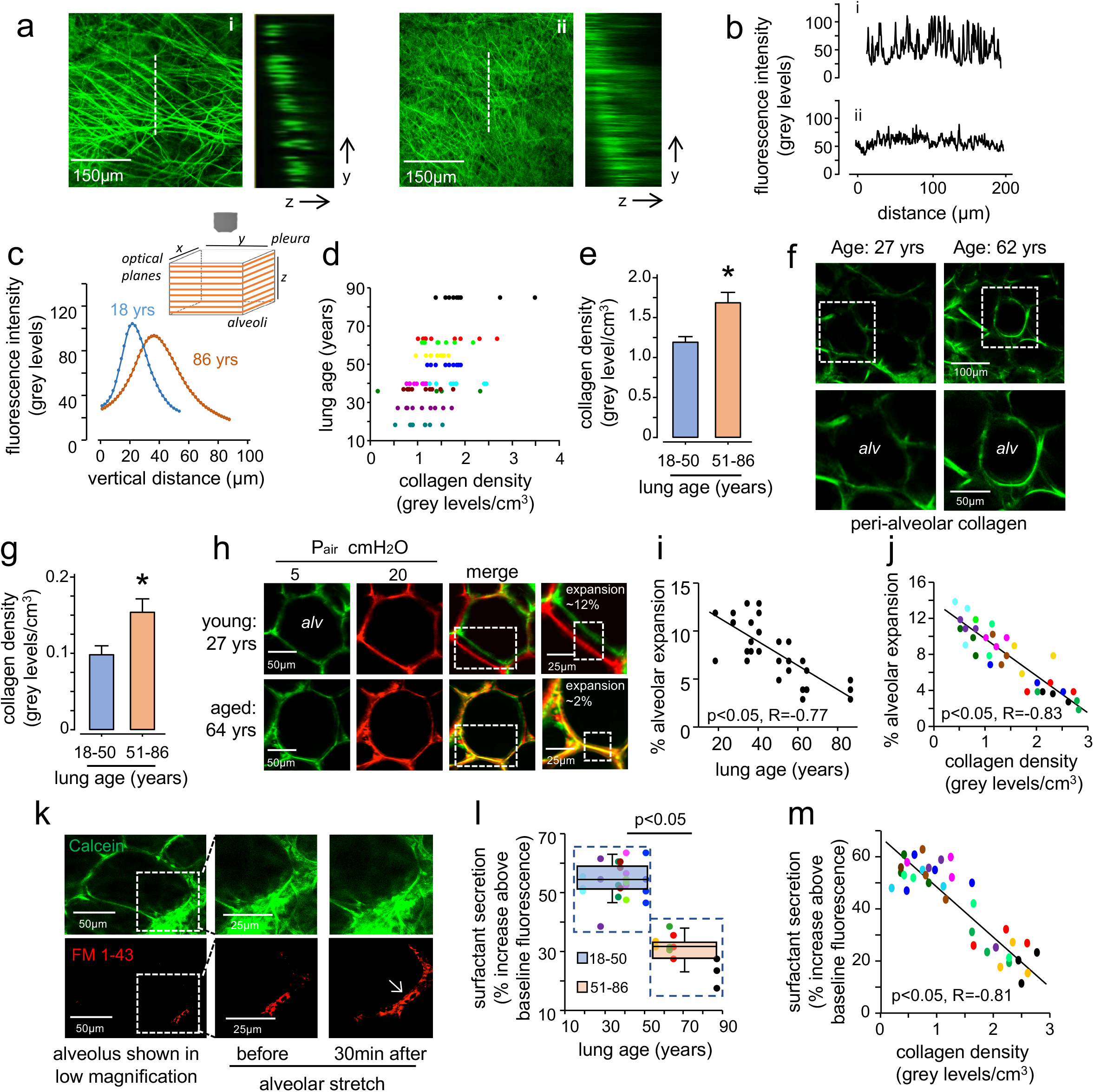
Aging-associated fibrosis limits alveolar expansion and surfactant secretion. **a-b** Two-photon images (*i and ii*) show collagen fluorescence by second harmonic generation in the subpleural interstitium of an 18 (left) and an 86 (right) year-old lung. Adjacent panels show collagen fluorescence in the depth plane (*y-z*) along the indicated lines (*dashed lines*). Tracings in (b) represent fluorescence intensity along the lengths of the dashed lines (“distance”). **c-e**, Imaging was carried out across a tissue volume calculated as the product of the area and the depth of the imaged field (see sketch). Tracings in (c) are from two representative fields and quantify collagen fluorescence along the depth axis from the pleural margin. In (d) “collagen density” was calculated as the summed fluorescence per unit volume for multiple fields in each lung. (Each color indicates a separate lung.) Group data are shown in (e). Bars: Mean±SEM, n=7 and 4 lungs respectively for 18-50 and 51-86 groups. ^∗^p<0.05 versus 18-50 group by Student’s t-test. **f-g**, Images and group data show peri-alveolar collagen. Bars: Mean±SEM, n=7 and 4 lungs respectively for 18-50 and 51-86 groups. ^∗^p<0.05 versus 18-50 group by Student’s t-test. **h**, Confocal images show an alveolus (*alv*) stained with the intracellular dye calcein-AM. Alveoli were imaged at alveolar pressures 5 (*green*) and 20 cmH20 (*red*). A select region (*rectangle in merge*) was magnified to show alveolar expansion during stretch. **i-j**, Alveolar expansions were plotted for individual imaged fields versus donor age (*i*) and subpleural collagen densities (*j*). P values were computed by linear regression. **k**, The images show an alveolus stained with calcein-AM (*green*), and the extracellular lipid dye, FM1-43 (*red*). A selected region (*rectangle in left image*) was magnified in the middle and right images. Alveolar stretch caused surfactant secretion as indicated by time-dependent increase of red fluorescence (*arrow*). **l**, Group data show stretch-induced surfactant secretion responses. Mean±SEM, n=7 and 4 lungs respectively for 18-50 and 51-86 groups. P value was computed by Student’s t-test. **m**, Surfactant secretion to alveolar stretch is plotted as a single dot for each alveolus across the indicated subpleural collagen density range. P value was computed by linear regression.

To determine whether increased collagen density impeded alveolar expansion, we carried out optical quantification of alveolar dimensions at low and high transpulmonary pressures (**Figure 4h**). This analysis demonstrated that there was a monotonic loss of alveolar expansion with age (**Figure 4i**) and that this loss correlated with collagen density (**Figure 4j**). Thus with age, alveolar expansion was limited by a constraining effect of increased peri-alveolar and subpleural collagen. Alveolar expansion causes secretion of surfactant, which maintains alveolar patency and provides epithelial defense against inhaled pathogens. To quantify the variability of surfactant secretion, a fluorescence approach was used quantify surfactant secretion at the single alveolar level.^32^ Our data indicate a strong negative correlation of surfactant secretion with age and with collagen density (**Figure 4k-m**).

Alveoli serve as essential gas exchange units that must inflate for effective ventilation, a process that deteriorates with age and can be limited by fibrosis^33^. Our results reveal cellular and molecular changes that occur in human lung aging. These include telomere shortening, loss of cellular proliferation, activation of cellular senescence programming, TGFβ signaling, and increased expression of collagen-regulatory genes. These changes lead to loss of alveolar type 2 cells, fibroblast expansion, and accumulation of interstitial collagen. Live lung imaging revealed regions of interstitial fibrosis in aged lungs, which were associated with local alveolar dysfunction. Collectively, these results shed light on the molecular pathways underlying fibrotic evolution in natural lung aging.

Acute respiratory distress syndrome (ARDS) is a major cause of mortality from acute lung infections and injury, and advanced age is associated with worse outcomes.^34^ Recent data have demonstrated an association between the subclinical fibrosis seen with aging, or interstitial lung abnormalities, and severe ARDS.^35^ Furthermore, telomere shortening in peripheral blood leukocytes was associated with increased severity and mortality from ARDS.^36^ The lung parenchymal telomere shortening observed in our study, the associated cellular senescence, and the resulting pro-fibrotic change suggest a mechanism for respiratory vulnerability with normal aging. Furthermore, the findings implicate a mechanism in common with pathologic lung fibrosis, where telomere shortening is a root cause^13,15^. Our study does not distinguish chronological aging from environmental insults accumulated over the lifespan, which are likely to be relevant given many genes and pathways in common with sun-exposed skin. Future studies should build on these findings and test the relative weight of cell autonomous and environmental effects, and also how senescence programs impair reparative responses to incident lung injury and infection.

## Methods

### Participants

RNA-seq, type 2 cell immunofluorescence, and telomere length analyses were done with the Lung Aging Cohort, which consists of 86 donor lungs collected between 2012 and 2018 and made available by the Donor West Network^37^. Fresh tissue fragments were snap-frozen in liquid nitrogen within 48 hours of x-clamp. Age, sex, ethnicity, smoking status, and cause of death were recorded. Second harmonic microscopy and surfactant studies were done with intact human lungs obtained from brain dead organ donors at the time of tissue acquisition for life-saving transplantation as described^38-40^ through a collaboration and protocol with LiveOnNY, the organ procurement organization for the New York area. Demographic data are detailed in **Table S1**.

### Bulk RNA Sequencing

Total RNA was isolated using the miRNeasy Mini Kit (Qiagen, Valencia, CA, USA). Extracted RNA samples were sent to Novogene for library construction and sequencing. Quantitation and quality control were done in three steps including NanoDrop (Thermo Fisher Scientific Inc., Waltham, MA), agarose gel electrophoresis, and Agilent 2100 Bioanalyzer (Agilent Technologies, Palo Alto, CA). mRNA was enriched using oligo(dT) beads and fragmented, and then cDNA was synthesized. Purified and processed cDNA libraries were checked on Agilent 2100 for insert size and quantified on Qubit and by qPCR. PE 150bp sequencing was done on Novaseq6000 machines to a sequencing depth of at least 6Gb for each sample. Adapter trimming and alignment to the reference genome were done using STAR software.^41^ Multi-factor differential expression analysis for age, smoking, and gender was done with DESeq2 and the likelihood ratio test was used for hypothesis testing.

### Telomere length qPCR

Genomic DNA was isolated from snap-frozen lung tissues using the Gentra Puregene Kit (Qiagen). DNA samples were run on 1% agarose gel electrophoresis for quality control and quantified using the NanoDrop spectrophotometer (Thermo Fisher Scientific Inc.). For each sample, cycle threshold values for telomere and the reference housekeeping gene (36B4) were determined in triplicates using quantitative PCR, as previously described^42,43^. Delta Ct was calculated by subtracting the mean telomere cycle threshold from the mean 36B4 cycle threshold. Three cell line standards with known telomere lengths were used to graph a standard curve, from which sample telomere lengths were calculated. Samples with standard deviation of triplicates higher than 0.25 were excluded.

### Ingenuity Pathways Analysis, Gene Set Enrichment Analysis, and ssGSEA

Pathway Analysis was done using the Ingenuity Pathways Analysis software (Qiagen). Differentially expressed genes with p ≤ 0.05 were used for analysis. Analysis results with p ≤ 0.05 were considered significant. Gene Set Enrichment Analysis was done on the GSEA software.^44,45^ Pearson metric was used for ranking genes, and a weighted enrichment statistic was used. Single sample gene set enrichment scores were computed on R using Singscore^46^.

### Analysis of publicly available data

The Genotype-Tissue Expression (GTEx) Project^10^ data (release V8) used for the analyses were obtained from the GTEx Portal (https://gtexportal.org/) on 8/20/2020. RNA-seq gene read counts, sample attributes, and subject phenotypes were downloaded for differential expression and subsequent analyses. Human lung single cell data used for cell type deconvolution analysis were downloaded from Reyfman et al^26^ (GSE122960).

### Immunohistochemistry and Immunofluorescence

Lungs were fixed with 10% formalin overnight and transferred to 70% ethanol before embedding in paraffin and sectioning to 4µm thickness. For immunohistochemistry of gamma-h2ax, sections were deparaffinized in xylene and rehydrated in an ethanol gradient series. Antigen retrieval was done by microwaving for 6 minutes in citrate buffer pH 6 (Sigma). After quenching in 3% H2O2 in Methanol, sections were permeabilized in 0.5% Triton X-100 in PBS. Sections were blocked in 3% BSA, 0.1% Triton X-100, 5% normal goat serum in PBS for 1 hour and incubated at 4°C overnight with primary antibody (Biolegend, cat. 613402, dilution 1:500), then at room temperature for 3 hours with secondary antibody (Santa Cruz Biotechnology, cat. sc-2005, 1:1000). Sections were developed in DAB working solution (Vector Laboratories) for 8 minutes, washed, dehydrated, and mounted with Cytoseal. All washes between steps were done with 1X PBS.

For immunofluorescence of pro-SPC, sections were deparaffinized, antigen retrieved, and permeabilized. Sections were blocked in 3% BSA, 0.1% Triton X-100, 5% normal donkey serum in PBS for 1 hour and incubated at 4°C overnight with primary antibody (EMD Millipore, cat. Ab3786, 1:300), then at room temperature for 1 hour with Alexa Fluor 594-conjugated secondary antibody (Life Technologies, cat. A21207, 1:1000). Sections were then washed and mounted with mounting medium with DAPI (Vector Laboratories). All washes between steps were done with 1X PBS on Day 1, and PBST (1:1000) on Day 2. Images were acquired on a Zeiss Axioscope 5 microscope. Immunoreactive cells were counted while blinded to the ages of the immunostained lungs.

### Cell type deconvolution

Cell type deconvolution of bulk RNA-seq data from the LAC was performed with MuSiC^27^, a publicly available computational resource. ScRNA-seq data from 3 young lungs aged 20-30 years (donors 3, 6, and 8) published by Reyfman et al.^26^ were clustered by Seurat^47^ and annotated for cell type by SingleR^25^ followed by the MuSiC workflow for cell type proportion analysis.

### Live two-photon imaging of human lungs

Two-photon and confocal microscopy were carried out on live, de-identified human lungs obtained after ∼20 hours of cold ischemia. The lingular lobe, which provides a flat surface convenient for live microscopy, was positioned below the objective of a two-photon microscope (TCS SP8, Leica). The lobe was perfused with buffer through the cannulated lobar artery at infusion pressure of 10 cmH2O, while inflated at alveolar pressure of 5 cmH2O through a bronchial cannula. We subpleurally injected fluorescent dyes through a 31-gauge needle. We detected subpleural collagen as the fluorescence of second harmonic generation (SHG) at an excitation and emission wavelengths of 830 nm and 425-460 nm, respectively. Non-specific autofluorescence and photobleaching were eliminated by appropriate gain setting. We quantified subpleural collagen density as the integrated collagen fluorescence per cubic centimeter in the space between the visceral pleura and the alveolar epithelium. Stretch-induced surfactant secretion was initiated by a single 15-second hyperinflation induced by increasing airway pressure from 5 to 15 cmH2O. We quantified surfactant secretion by the timed appearance of lipid-sensitive fluorescence in the alveolar space^32^.

### Statistical Analysis

Statistical analysis for comparison of two groups was done using the unpaired, two-sided, two-samples t-test. For comparison of multiple groups, 1-way ANOVA was used. Pearson correlation coefficient R was calculated to assess association of two continuous variables. Unless otherwise stated, a p-value less than 0.05 was considered significant. Multiple hypothesis testing using Benjamini-Hochberg method was done when appropriate.

### Study Approval

Tissue samples were obtained from brain-dead (deceased) individuals, and thus this study does not qualify as human subjects research, as confirmed by the UCSF and Columbia University IRBs.

## Supporting information

Lung donor demographic data

Ingenuity Pathways Analysis

Genes differentially expressed by human lung epithelial cells from GSE122960

Genes differentially expressed by human lung fibroblasts from GSE122960

## Author contributions

J.L. did telomere length measurement and immunofluorescence and performed computational analyses with help from K.B. and L.M. and under the supervision of M.B., D.A., W.E., and S.C. D.J.D. and J.R.A. derived and helped J.L. in applying the consensus senescence marker analysis. R.M. and M.K. procured LiveOnNY donor network lungs under the supervision of D.L.F. M.N.I. performed second harmonic and surfactant microscopy under the supervision of J.B., S.B., and D.L.F. M.B., P.J.W, and J.B. conceived of the work, supervised experimental planning and analysis, and co-wrote the manuscript with input from M.M.

## Acknowledgements

This work was supported by NIH HL131560 and institutional startup funds to M.B., NIH HL139897 to P.W., HL145547 to D.F. and J.B., NIH HL36024 and HL57556 to J.B., and the Nina Ireland Program for Lung Health to M.B. and P.W.

## Competing Interests

D.J.D. and J.R.A. are current employees of Genentech and shareholders in Roche.

