## Supplementary Figure S1 for "Molecular programs of fibrotic change in aging human lung"

**<u>Supplementary Figure 1</u> a**, Cell type deconvolution for LAC and Gtex Lung. Age for GTex is represented in decades. Pearson R and and p values are shown for LAC, and 1-way ANOVA is shown for GTex. **b**, List of collagen processing and cross-linking genes used for Figure 3c (bolded genes were enriched with age in the lung aging cohort). **c**, Single sample gene set enrichment analysis for the collagen processing genes enriched in aging from in the oldest and youngest quintiles from multiple GTex tissues. P values are for two-tailed Student's t-test.

a

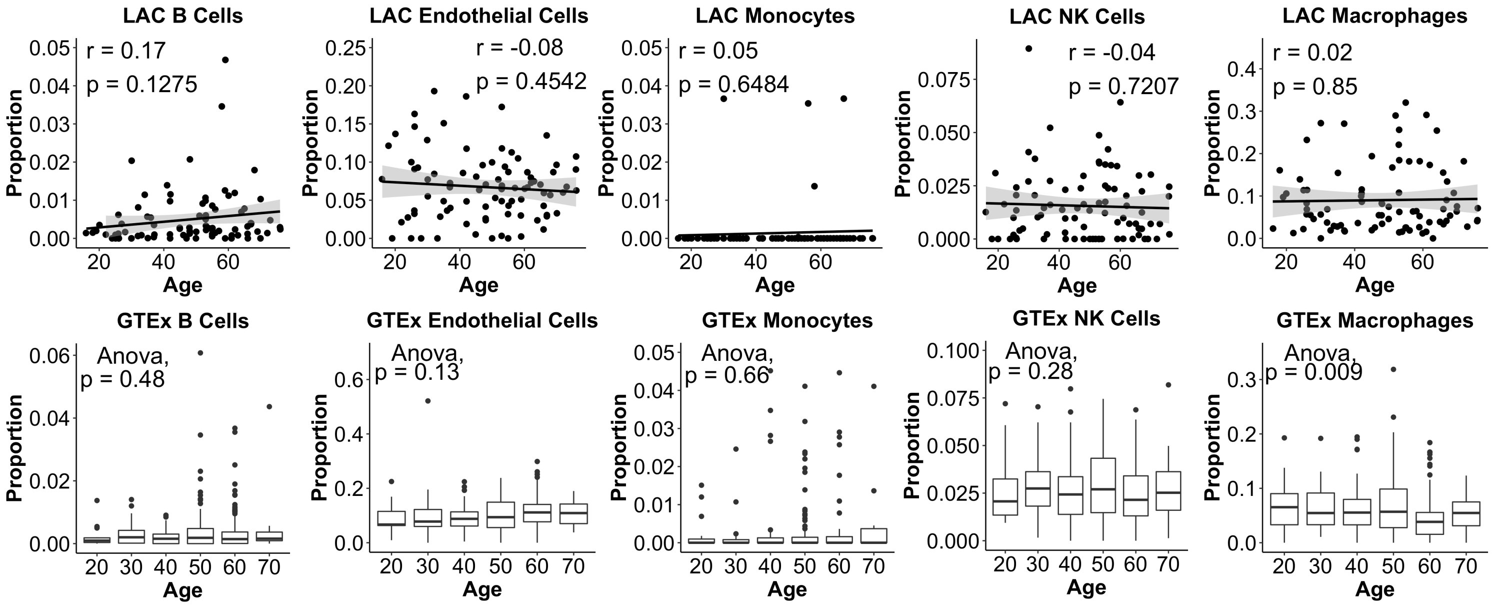


b

COLGALT1
**COLGALT2**
**LOX**
**LOXL1**
LOXL2
**LOXL3**
**LOXL4**
**PCOLCE**
**PCOLCE2**
TGM1
**TGM2**
**TGM3**
TGM4
**TGM5**
TGM6
TIMP1
**TIMP2**
**TIMP3**
TIMP4

c

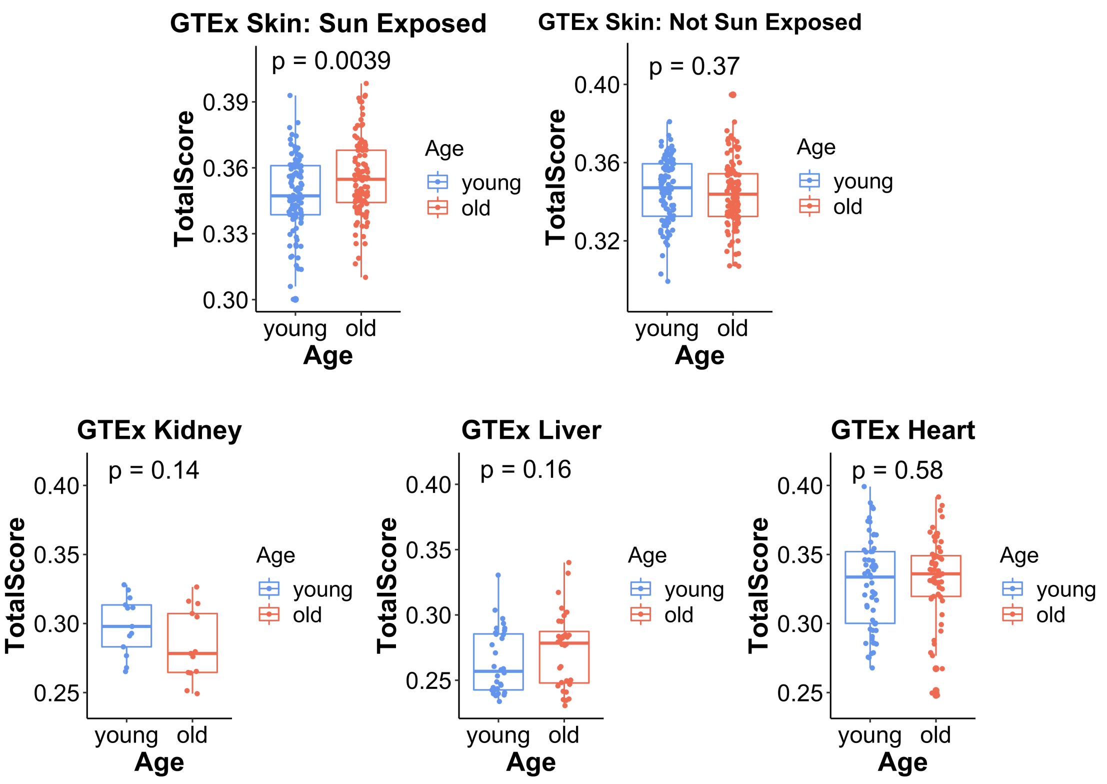
